# Highly and lowly domesticated endangered fish from a conservation hatchery diverge in their thermal physiology, transcriptome, and methylome

**DOI:** 10.1101/2025.08.08.669364

**Authors:** Joanna S. Griffiths, Amanda J. Finger, Melinda R. Baerwald, Md Moshiur Rahman, Tien-Chieh Hung, Nann A. Fangue, Andrew Whitehead

**Affiliations:** Department of Environmental Toxicology, University of California Davis, Davis, CA 95616; Conservation Biology Division, Northwest Fisheries Science Center, National Marine Fisheries Service, National Oceanic and Atmospheric Administration, Seattle, WA, USA; Department of Animal Science, University of California Davis, Davis, CA 95616; Division of Integrated Science and Engineering, California Department of Water Resources, Sacramento, CA 95814; Department of Biological and Agricultural Engineering, Fish Conservation and Culture Laboratory, University of California Davis, Davis, CA 95616; Department of Wildlife Fisheries and Conservation Biology, University of California Davis, Davis, CA 95616

**Keywords:** phenotypic plasticity, developmental plasticity, domestication selection, critical thermal maximum, ecophysiology, conservation physiology

## Abstract

Conservation hatcheries aim to produce fish for supplementation of wild populations, but hatchery environments may drive phenotypic divergence from wild fish. These diverged traits may have reduced fitness in the wild, which could compromise wild population sustainability and evolutionary potential, such as in response to climate change. Delta smelt are a critically endangered fish species that are safeguarded against extinction with a hatchery refuge population. We investigated whether elevated rearing temperature through larval development adjusted upper thermal tolerance limits (acclimation) in Delta smelt, whether upper thermal tolerance and plasticity (acclimation ability) differed between fish with old or recent hatchery ancestry (high or low domestication index; DI), and temperature and DI effects on liver transcriptome and methylome patterns. We observed that elevated rearing temperatures induced higher thermal tolerance (acclimation). Individuals with higher DI also had higher upper thermal tolerances, but high DI families had reduced thermal plasticity between rearing temperatures. This is consistent with domestication causing heritable elevation of upper thermal tolerance but at the cost of reduced thermal plasticity. High and low DI fish were differentiated in both genetic variation and methylome variation, suggesting the influence of both during domestication. But methylome differences distinguishing high and low DI fish did not overlap with temperature-induced methylome changes, and do not appear to be stably inherited in the hatchery. We conclude that domestication selection has altered thermal physiology within the refuge hatchery despite careful genetic management, underpinned by shifts in the transcriptome and methylome. These changes could affect Delta smelt fitness upon reintroduction to habitats that continue to warm, and show that physiological traits can diverge even within carefully genetically managed hatchery populations.

## Introduction

A major goal of captive breeding programs and conservation hatcheries is to preserve genetic diversity and evolutionary potential for at-risk species (Fisch et al. 2015; Russello and Jensen 2018). However, captive rearing can cause unintended genetic consequences, such as genetic drift, inbreeding, and adaptation to captivity (domestication selection). As a result, phenotypic and genetic divergence may occur between wild and hatchery individuals, often with hatchery individuals maladapted to wild conditions (Laikre et al. 2010; Lorenzen et al. 2012). Maladaptation upon reintroduction to the wild may be exacerbated if natural environments are rapidly changing such that optimum phenotypes are even further diverged from those favored in hatcheries (e.g., rising temperatures due to climate change; Tillotson et al. 2019; Austin et al. 2021; Wilder et al. 2024). Fish reared in static hatchery environments that are designed to provide optimal conditions for growth and reproduction may be unprepared for the fitness challenges faced in natural environments, especially given the pace of environmental change in the Anthropocene (Mason et al. 2013; Milla et al. 2021; LaCava et al. 2023). Identifying phenotypic divergence in captivity, including the drivers and underlying molecular mechanisms, will illuminate the multi-generational impacts of captivity, the potential impacts of captivity on future persistence of species upon release in changing environments, and may inform changes in captive environments to enhance the resilience of endangered species.

Within hatcheries some phenotypes may be favored over others, with variance inherited through either genetic or non-genetic (epigenetic) mechanisms. Inheritance of environmentally induced epigenetic changes, including histone modifications or DNA methylation (Burggren 2016), is an increasingly appreciated mechanism to explain phenotypic divergence between hatchery and wild fish, where changes can emerge in just one or few generations. Most environmentally induced epigenetic changes are reset each generation, but occasionally some may be stably inherited. A single generation of hatchery rearing has caused epigenetic divergence from wild counterparts in fish including Nile tilapia (Konstantinidis et al. 2020), Pacific Salmon (Le Luyer et al. 2017), and Delta smelt (Konstantinidis et al. 2020; Habibi et al. 2024). Hatchery rearing may induce epigenetic change that alters gene expression (Konstantinidis et al. 2020) which may facilitate phenotypic change (Jaenisch and Bird 2003) that distinguishes hatchery-reared fish from wild counterparts. Phenotypic variation favored in hatchery environments may also be underlain by genetic variation (i.e., domestication selection) (Howe et al. 2024). It is difficult to attribute genetic or epigenetic causes for hatchery-induced phenotypic change, but this may be important since these mechanisms may differ in their pace of change, the reversibility of change, and in the types of management practices that could prevent change.

Delta smelt (*Hypomesus transpacificus*) is a critically endangered osmerid fish (Finger et al. 2018; California Natural Diversity Database (CNDDB) 2024; Habibi et al. 2024), that is endemic to the Sacramento-San Joaquin Delta (Delta) and is supported by a conservation hatchery. Wild populations are threatened by many factors, including rising temperatures and salinities, competition with non-native species, exposure to environmental pollution, and water management conflicts (IEP MAST 2015; FLOAT-MAST and (Flow Alteration - Management Analysis and Synthesis Team) 2022). A captive breeding program was initiated at the University of California Davis Fish Conservation and Culture Laboratory (FCCL) in 2008 where the species is genetically managed in the hatchery with deliberate breeding that uses genetic and pedigree information to avoid mating of relatives, equalize family sizes, and maintain genetic diversity by introducing wild fish each year, depending on availability (Lindberg et al. 2013). Despite intensive genetically informed management, phenotypic differences that distinguish hatchery from wild fish have been emerging, including increased reproductive success and age at maturity in the hatchery for fish with deeper hatchery ancestry (Finger et al. 2018; Ellison et al. 2023; LaCava et al. 2023; Tsai et al. 2023). A possible mechanism of phenotypic change in captivity could be genetic change (e.g., captivity-induced selection), but there is also evidence for environmentally induced change (phenotypic plasticity) including hatchery-induced epigenetic changes that occur within the first generation (Habibi et al. 2024). Heritable hatchery-induced phenotypic changes could disadvantage hatchery fish in the wild, as has been observed in salmon (Laikre et al. 2010; Lorenzen et al. 2012). Furthermore, hatchery-induced phenotypes may compromise fitness not only in contemporary wild environments, but also in near-future environments that are rapidly changing because of climate change (Tillotson et al. 2019; Austin et al. 2021). On the other hand, there is evidence that domesticated fish show lower stress levels (due to handling in the hatchery environment) and higher environmental tolerance, such as hypoxia tolerance, which may impart higher fitness in the wild (Scott et al. 2014; Ghazal et al. 2025). Consequently, there is a need to understand how domestication selection and hatchery rearing in Delta smelt has altered phenotypes that are important in contemporary natural environments and future environments that are rapidly changing.

Resilience to elevated temperature and temperature extremes are a key concern for the conservation of wild species in the face of climate change. Delta smelt are particularly sensitive to elevated temperature compared to other native and non-native species in their watershed (Swanson et al. 2000; Davis et al. 2019a; Hung et al. 2022). Projected temperature increases in the San Francisco estuary due to climate warming indicates that physiological limits for Delta smelt have already been exceeded, thereby threatening fitness (Cloern et al. 2011; Komoroske et al. 2014; Brown et al. 2016; Jeffries et al. 2016). Thermal plasticity (acclimation ability) is one way for fish to increase their upper thermal tolerance limits. Hatchery practices that promote tolerance to ecological stressors, such as through warm acclimation, may enhance performance and survival when released into the wild. However, acute thermal acclimation in Delta smelt is accompanied with a physiological stress response, such as the upregulation of cellular stress response genes (Komoroske et al. 2015; Jeffries et al. 2016), and changes to behavior and predator avoidance (Davis et al. 2019b) suggesting potential costs or tradeoffs to warm acclimation. Importantly, it is unknown whether hatchery domestication has altered Delta smelt’s thermal physiology and plasticity.

We designed experiments to test whether rearing temperature in the hatchery affected thermal acclimation, and whether acclimation capacity differed depending on domestication history. First, we examined the extent to which Delta smelt can adjust their upper thermal tolerance limits through acclimation by exposure to ecologically relevant rearing temperatures (15℃ and 18℃) and characterized underlying molecular changes (transcriptome and methylome). We then tested whether thermal tolerance limits differ between fish with low levels of hatchery ancestry compared to those with extensive hatchery ancestry and is there transcriptomic and epigenomic divergence between low and high domestication fish? Delta smelt routinely experience temperatures between 15 and 23 ℃ in the wild (Jeffries et al. 2016) and these temperatures have continued to increase in recent years (Bashevkin et al. 2022). In contrast, Delta smelt in the hatchery are reared at a constant 16℃, which may result in different baseline tolerances. Finally, we test whether low and high domesticated Delta smelt have diverged plasticity (acclimation abilities) in their ability to change their upper thermal tolerance, transcriptome, and methylome in response to elevated rearing temperatures?

## Methods

### Adult spawning and offspring rearing

Our source of Delta smelt was the refuge population at FCCL which has been developed and maintained since 2008. We examined fish with different degrees of domestication, where the extent of hatchery ancestry is defined as the fish’s domestication index (DI). DI is an additive metric that measures the number of generations an individual’s genome has spent in captivity. Wild fish, when available, were introduced each year and bred with a hatchery individual. Wild fish have a DI of 0 and offspring from wild parents have a DI of 1. When parents have different DIs, the DI of the offspring is the average DI of the parents plus 1. In our experiment, parent DIs spanned a range from 5 to 10. In 2021, we spawned adult Delta smelt following a modified North Carolina II breeding block design (Figure S1). We crossed 15 males of high DI with 6 females of high DI and 15 males of low DI with 6 females of low DI. Low DI families are defined as a DI <7, and high DI families have a DI >9. We were unable to include individuals with lower DI since wild individuals have not been available for breeding in recent years (Hung 2025). For each experimental breeding block (total of 12 blocks), eggs from two individual females were independently crossed with sperm from five individual sires for a total of ten separate fertilizations.

At 3 days post fertilization (dpf), offspring from each family were split evenly for rearing in one of two temperature treatments, including two replicates at 15℃ and two replicates at 18℃ (Figure S1), in two parallel recirculating aquaculture systems (Tsai et al. 2022). Families were assigned to two different DI categories (low or high) and reared separately at either 15℃ or 18℃ (two DI categories at two rearing temperatures each with two duplicates, equals 8 chambers total). Per family a minimum and maximum of 10 and 60 embryos were added per chamber. Families with insufficient embryos to reach the minimum per-treatment distribution were excluded. Hatching occurred after ∼10 dpf and larvae were subsequently moved to 10-gallon trays and at 40 dpf, larvae were moved into 133-L tanks, where they were reared up to 143-151 dpf for the completion of all experiments. Fish were held under natural photoperiod, salinity of 2.3 ppt, and fed live prey from cultures of rotifers (*Brachionus plicatus*) and brine shrimp nauplii (*Artemia franciscana*) (Lindberg et al. 2013). All procedures were approved by the UC Davis Institutional Animal Care and Use Committee (IACUC Protocol #21915).

### Thermal tolerance measurements

We determined upper thermal tolerance using a critical thermal maximum (CTMax) approach (Beitinger 2000) for larvae that were 71-79 dpf. This age was chosen to ensure we had high survival for large sample sizes in critical thermal methodology. Thermal tolerance trials were completed for 195 fish from each tank (n = ∼390 per treatment; 1560 fish total). We measured the thermal tolerance of 26 fish per trial. Fish were individually placed into 30-mL opaque cups with a gentle air bubbler for aeration and to prevent thermal stratification. Cups were placed into temperature controlled water baths, where portable heaters steadily increased the temperature of the cups by 0.3℃ min^-1^ until fish experienced loss of equilibrium (LOE), a common endpoint used to determine thermal tolerance limits (Beitinger et al. 2000; Komoroske et al. 2014; Jeffries et al. 2016; Davis et al. 2019a). Cup water temperature at LOE was recorded with a calibrated immersion thermometer. Due to logistical constraints and the large number of fish that we measured, we were unable to recover our fish post CTMax measurements. Therefore, we performed a set of three preliminary trials on 63 fish to test our methods for potential biases. We determined that thermal tolerance measurements were not influenced by observer bias (F5,57 = 1.36 p = 0.25), suggesting that we had successfully chosen a repeatable endpoint. However, we observed high mortality 24 hours post recovery (89%), but higher thermal tolerance was not correlated with mortality (F2,60 = 0.55, p = 0.58). Because of this high mortality, we performed a control trial with 25 fish using the same methodology and fish handling as experimental trials, but there was no increase in temperature in the water bath. We found that mortality was at 40.9 % 24 hours post recovery. Overall, we conclude that due to the delicate nature of Delta smelt larvae, some mortality was due to handling during the experiment.

A linear mixed-effects ANCOVA (R package lme4 v1.1-35.1 (Bates et al. 2014) and car v3.1-2 (Fox and Weisberg 2019), was used to test for the fixed effects of domestication index (low or high) and rearing temperature (15° or 18°C), and their interaction, on CTMax (the response variable), where tank was nested within the recirculating system as a random effect (Figure S1). We included fork length as a fixed covariate since it had a significant correlation with CTMax for smaller fish (< 15mm). To meet the assumptions of the ANOVA, we performed an orderNorm transformation (bestNormalize v1.9.1 (Peterson 2021)) on CTMax.

We further investigated the plasticity of thermal tolerance by calculating mean CTMax at the family level (see methods below on “*Family-Level Assignment*” for identifying families) for both rearing temperatures, and then extrapolating the slopes (i.e., the difference in mean CTMax between 18℃ and 15℃for each family). We included all families with valid CTMax estimates at both rearing temperatures and each replicate (100 out of 121 families). We performed a linear model ANOVA (R package car v3.1-2 (Fox and Weisberg 2019)), with domestication index (low or high) as the fixed effect and the slope as the response variable. We repeated the linear model with higher precision by requiring at least 2 individuals per family per treatment (78 out of 121 families). We confirmed that mean family CTMax measurements were not biased by survival differences survival between fish of low or high DI fish (Figure S2).

### DNA and RNA extraction and library preparation for methylome and transcriptome sequencing

Fish were maintained at the two rearing temperatures until they were juveniles (143-151 dpf) and then sampled from the high and low DI groups for transcriptome and methylome analyses. At this age, juvenile fish were large enough to isolate and dissect the liver. Immediately following incapacitation (buffered MS-222), livers were dissected and flash frozen in liquid nitrogen and archived at −80° C. We chose liver because of its role in energy storage and metabolism (Harper and Wolf 2009). We collected 8-17 individuals from each treatment (temperature by DI by replicate) for a total of 102 individuals (only 3 individuals were available at the 18℃ Low DI replicate). For 40 of these samples (4-5 individuals per temperature per replicate per DI), we performed a dual DNA and RNA extraction from a single liver sample using the Qiagen AllPrep DNA/RNA Micro Kit (Valencia, CA). For the remaining 62 samples, we extracted DNA from frozen livers using the Qiagen DNeasy Blood & Tissue Kit (Valencia, CA).

We constructed 40 individually-indexed RNASeq libraries in a single batch using the NEBNext Ultra II Directional RNA Library Prep Kit (Illumina, San Diego, CA). We multiplexed all 40 RNASeq libraries into a single pool that was sequenced across two lanes of an Illumina HiSeq4000 (PE-150 bp; Illumina, San Diego, CA). Sequencing was performed by IDSeq Inc. (Davis, CA).

From the 102 samples for whole genome bisulfite sequencing (WGBS) analysis, 4pg of unmethylated lambda control DNA (Promega, Madison, WI) was spiked into 4.5ng genomic DNA for each sample, then bisulfite converted using the Zymo EZ Methylation Gold kit (Zymo Research, Irvine, CA) according to the recommendations of the manufacturer. Bisulfite-converted DNA was converted into individually-indexed (barcoded) sequencing libraries using the xGen Methylation-Sequencing DNA Library Preparation Kit (IDT, Coralville, IA) in a single batch. The fragment size distribution of the resulting libraries was assessed on the LabChip GX Touch system (PerkinElmer, Waltham, MA), quantified on a Qubit instrument (LifeTechnologies, Carlsbad, CA), and pooled in equimolar ratios. The library pool was quantified by qPCR with a Kapa Library-Quant kit (Kapa Biosystems/Roche, Basel Switzerland) and sequenced across two lanes of an Illumina Novaseq 6000 system (PE-150 bp; Illumina, San Diego, CA) at the UC Davis Genome Center.

### RNASeq and differential gene expression analysis

We developed a custom snakemake workflow for RNAseq quality control and transcript quantification (https://github.com/JoannaGriffiths/RNASeq-snakemake-pipeline), which included FastQC (v0.11.9 (Andrews 2010)) for quality control, fastp (v0.20.1 (Chen et al. 2018)) for trimming of adapters, and Salmon (v1.10.1 (Patro et al. 2017)) for transcript quantification and generation of a gene count matrix normalized by transcripts per million (TPM). RNAseq reads were aligned to the *H. transpacificus* transcriptome assembly (mapping-based mode of Salmon) and we used the reference genome assembly (GCF_021917145.1) to create a file of decoy-aware sequences. The read count matrix was filtered to retain genes with a minimum expression threshold of at least 0.5 TPM from all 8 replicate individuals from at least one treatment (i.e., domestication by temperature).

To visualize experiment-wide transcriptome variation, we performed a principal coordinate analysis (PCoA) using Euclidian distances on the log-transformed filtered and normalized RNAseq data set using the R package vegan v2.6-4 (Oksanen 2015). From the PCoA, we discovered and removed an outlier (a low DI individual from the 18°C exposure treatment), followed by another round of filtering and normalization. Differential gene expression analysis was performed using the R package limma v3.58.1 (Ritchie et al. 2015). We tested for the influence of two main effects of temperature and domestication index and their interaction, with replicate tank treated as a random effect. We used the global fit method in limma-voom and false-discovery rate (FDR) correction of p-values. Heatmaps were created with the R package gplots v3.13.1 (Warnes et al. 2016). Gene Ontology (GO) enrichment was tested using a Fisher’s exact test for up and down regulated genes separately (Wright et al. 2015), where the background set was the set of filtered genes used for discovery of differential gene expression. Significant functional enrichment included FDR adjustment of p-values and required the GO category to contain at least 5 genes. GO terms were identified by matching Delta smelt gene names with those from the zebrafish annotation database on ensembl using the R package biomaRt (Durinck et al. 2005, 2009).

### Bisulfite mapping, CpG identification, and differential methylation analysis

For WGBS samples, we used the program FastQC v0.11.9 (Andrews 2010) for quality control and removed four samples that had either high PCR duplication or were outliers in PCA (three high DI individuals from the 18°C exposure treatment and a high DI individual from the 15°C exposure treatment). TrimGalore (v 0.6.6, https://www.bioinformatics.babraham.ac.uk/projects/trim_galore/) was used to trim adapters and remove poor quality bases (Q < 30). Reads were mapped to the *H. transpacificus* reference genome assembly (GCF_021917145.1) using Bismark (v0.22.3, (Krueger and Andrews 2011) and bowtie2 (v2.4.2). PCR duplicates were marked using Bismark (v 0.22.3). We quantified genome-wide coverage using Qualimap (v 2.2.1 (Okonechnikov et al. 2016)). We used Bismark to call methylation sites and generate a CpG report as input for differential methylation analysis.

We used DMRichR (v.1.7.8, (Laufer et al. 2020)) for statistical analysis and visualization of differentially methylated regions (DMRs). We used a generalized least squares (GLS) regression model with permutation testing, where filtration for a region to be considered differentially methylated required a minimum of 5 CpG sites, coverage of 1x in at least 75% of samples, and a cut-off of 0.05 for the CpG coefficient used to discover testable background regions. We ran two independent models (for 10 permutations each), with rearing temperature or domestication index as the test covariate for each model. Random effects were not supported in DMRichR, so replicate tank was modeled as a covariate. Sex was unknown for individuals and was therefore not included as a covariate.

DMRs and background datasets (generated from DMRichR) were annotated using HOMER (v4.11.1 (Heinz et al. 2010)) and the genome annotation file from NCBI (GCF_021917145.1). HOMER uses the annotatePeaks.pl script to find the nearest transcription start site for each DMR (default parameters restricted to 1kb upstream and 100bp downstream). HOMER annotation also performs a relative enrichment of DMRs in each set of genomic annotations using the binomial distribution to assign a significance p-value. GO enrichment among DMRs was tested using a Fisher’s exact test for hyper- and hypo-methylated DMRs separately (Wright et al. 2015), with the background set generated from the DMRichR analysis. We first tested for functional enrichment using DMRs from annotation categories defined by HOMER software (promoters, exons, and introns), then we tested only DMRs that overlapped with promoter regions, and finally we tested for DMRs that overlapped gene bodies (exons and introns). Determination of GO enrichment used the same criteria as for transcriptomics.

We further investigated the functions of DMRs due to DI by comparing overlap with temperature-induced DMRs. If there was a significant number of DMRs common between DMRs due to DI and rearing temperature, this would suggest that DMRs due to DI are regions of the genome that may also be important for elevated temperature tolerance. We used a hypergeometric test (phyper function in base R v4.3.2) to determine whether the significant overlap was more than expected by chance given the overlapping background set of 1,797 genes in both datasets.

We also sought to test whether the methylation response to rearing temperature differed between high and low DI fish (temperature-by-DI interaction). DMRichR software does not permit formal testing for interactions between two factors. We therefore adopted a conservative alternative approach to identify temperature-by-DI interactions; we tested for temperature treatment effects in high DI fish and low DI fish separately, then for temperature-induced DMRs that were significant in both high and low DI fish we tested whether temperature effects were in the same or opposite directions (e.g., hypo- vs. hyper-methylated) between DI groups. Cases where the direction of differential methylation was opposite between DI groups were considered indicative of temperature-by-DI interactions.

Environmentally induced methylation was observed in the first few hatchery generations of Delta smelt (Habibi et al. 2024). We intersected DMRs from our results with overlapping DMRs from Habibi et al. (2024) to test: 1) whether temperature induced DMRs (our data) were also differentially methylated during the earliest hatchery generations (Habibi et al. 2024 data); and 2) whether DMRs during the earliest hatchery generations (DMRs from Habibi et al. 2024) were further differentially methylated in later hatchery generations (our DI-induced DMRs). Although the two studies examined different tissues, both had a similar sequencing depth (10x), sample size (∼100 individuals) and the same bioinformatic pipeline for statistical analyses. Therefore, we considered results at least qualitatively comparable, especially if one were to expect that stable inheritance of epigenetic modifications would manifest in similar ways across multiple tissues.

### DMR and DEG correlation

We tested for genes that were both significantly differentially methylated and differentially expressed for the 39 individuals that had both RNASeq and WGBS data. We used a hypergeometric test (phyper function in base R v4.3.2) to determine whether the significant overlap was more than expected by chance given there were 1,302 genes overlapping in both background datasets. For each overlapping gene, we ran a linear model in R testing the correlation between percent DMR and the normalized read count expression data. To account for multiple testing, we applied a 0.05 p-value Bonferroni correction (based on the number of overlapping genes that were differentially methylated and differentially expressed) to determine the number of genes that showed a significant correlation.

### Family-level assignment and genetic differentiation between high and low DI fish

To assign offspring to families and estimate genetic differentiation, we sequenced whole genomes in 31 (774) low-DI adult progenitor (offspring) fish and 29 (776) high-DI adult progenitor (offspring) fish. DNA was extracted from fin clips as described in (Ali et al. 2016). Library preparation used the Twist 96-Plex Library Preparation protocol (Twist Bioscience, San Francisco, CA). We sequenced at a target depth of 30x coverage per progenitor and at 1-2x coverage per offspring (NovaSeq S4, PE-150 bp; Illumina, San Diego, CA). Using the snpArcher pipeline (https://snparcher.readthedocs.io/en/latest/index.html), reads were mapped to the *H. transpacificus* genome (GCF_021917145.1) and then genotyped with GATK (Van der Auwera & O’Connor 2020). Filters applied in GATK included: QD < 2.0, FS > 60.0, MQ < 40.0, MQRankSum < −12.5, ReadPosRankSum < −8.0, SOR > 3.0, GQ < 20. Additional filters removed SNPs with DP <2 and DP > 100 using vcftools (v1.14). We estimates a Weir pairwise F_ST_ (window size of 10,000 bp and a sliding window of 5000 bp) between low and high DI progenitors using vcftools (v0.1.16 (Danecek et al. 2011)). We ran a permutation test to determine whether F_ST_ was higher than expected by chance by randomly shuffled individuals into two groups (1,000 random permutations). Offspring genotypes were imputed (GLIMPSE v1.1.1 (Rubinacci et al. 2021)) using a phased reference panel of the progenitors (SHAPEIT5 v5.1.1; (Hofmeister et al. 2023)). A family-level pedigree was generated with progenitor and offspring genotypes using AlphaAssign (Whalen et al. 2019)

## Results

### Thermal tolerance measurements

There was a significant effect of rearing temperature on CTMax (F_1,5.0_ = 18.6 p= 0.008; Figure 1A). Mean CTMax of fish reared at 18°C was 0.63°C higher than fish reared at 15°C. There was a significant effect of DI on CTMax (F_1,4.0_ = 25.2 p = 0.007; Figure 1). Mean CTMax was 1.1°C higher for high compared to low DI fish. There was no interaction between temperature and DI (F_1,4.0_ = 0.001 p = 0.97; Figure 1). However, we did observe variation in plasticity at the family level when comparing their mean CTMax at 15°C and 18°C (Figure 1B). Additionally, high DI fish had reduced plasticity (slopes) in their response to rearing temperature compared to low DI fish (F_1,9_ = 5.5 p = 0.02; all families included; Figure 1C). This result remained significant when we required at least 2 individuals per family in each treatment (F_1,3.7_ = 4.3 p= 0.045).

**Figure 1:**
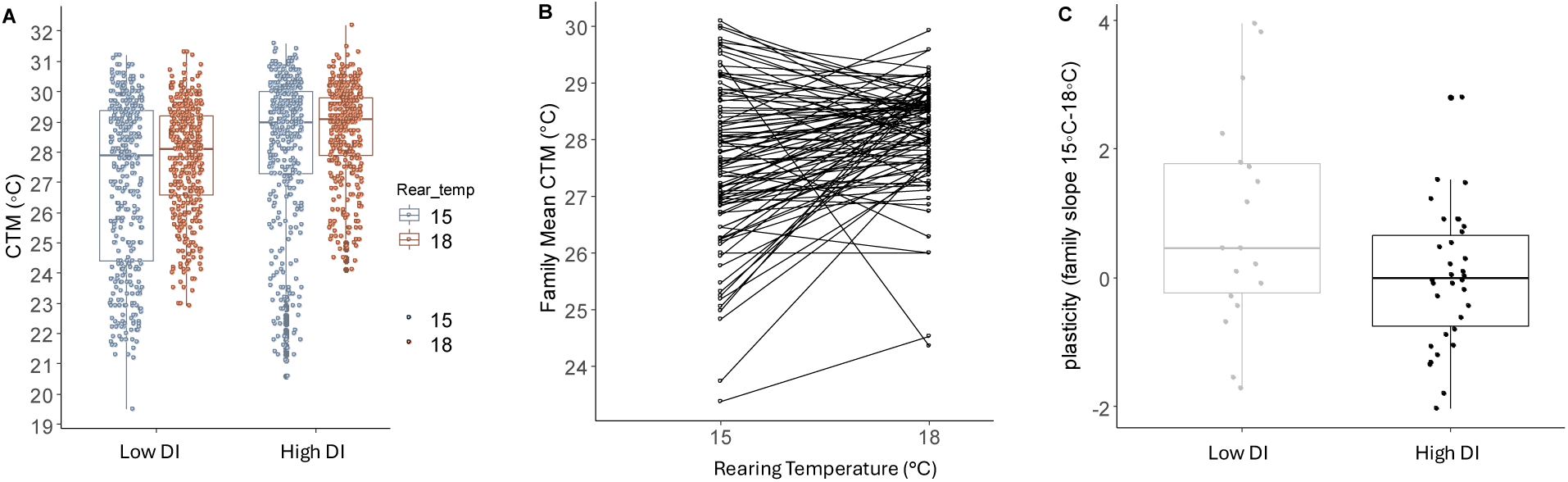
Critical Thermal Maximum (CTMax) variation within and among domestication index (DI), rearing temperatures, and families. A) CTMax for fish raised at either 15° C (blue) or 18° C (orange). Fish reared at 18° C had a higher CTMax than fish reared at 15° C (p=0.008) and high DI fish had higher CTMax than low DI fish (p=0.007). B) Variation in mean CTMax families (connected by black line) reared at either 15°C or 18°C. C) High DI families had reduced plasticity (the slopes of family means from Figure 1B) compared to low DI families (p=0.02).

### Differentially expressed genes

After normalization and filtration, we performed a PCoA to visualize liver transcriptome variation across all samples. PC1 and PC2 accounted for 23.02% and 10.35% of variation, respectively, with the effects of temperature appearing influential along PC1 (Figure 2).

**Figure 2.**
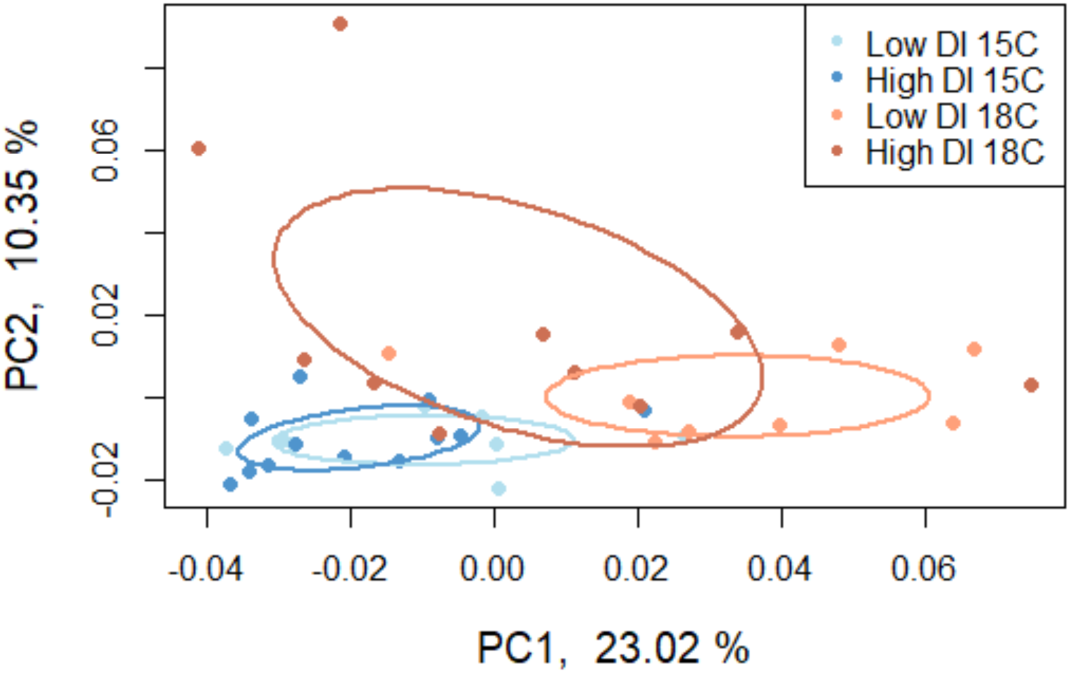
Euclidian distance-based principal coordinate analysis (PCoA) of read counts for all genes and individuals. Blue and orange colors represent samples from 15°C and 18°C treatments, respectively. Light and dark shades represent samples from low and high DI treatments, respectively.

We detected 1,609 differentially expressed genes (DEGs) between rearing temperatures with 683 genes more highly expressed at 18°C and 926 genes more lowly expressed at 18°C (Figure 3A). DEGs that were more lowly expressed at 18°C (Set 1) were enriched for Biological Processes GO terms including *small molecule metabolic process*, *small molecule catabolic process*, *sterol biosynthetic process, lipid metabolic process,* and *lipid biosynthetic process* (Figure 3B), and for Molecular Function GO terms including *oxidoreductase, isomerase, and insulin-like growth factor binding* (Figure 3C). DEGs that were more highly expressed at 18°C (Set 2) were enriched for Biological Processes GO terms including *response to stimulus*, *defense response*, *activation of immune response,* and *positive regulation of immune system process* (Figure 3D), and for Molecular Function GO including *peptidase activator, peptidase regulator,* and *immune receptor* (Figure 3E).

**Figure 3.**
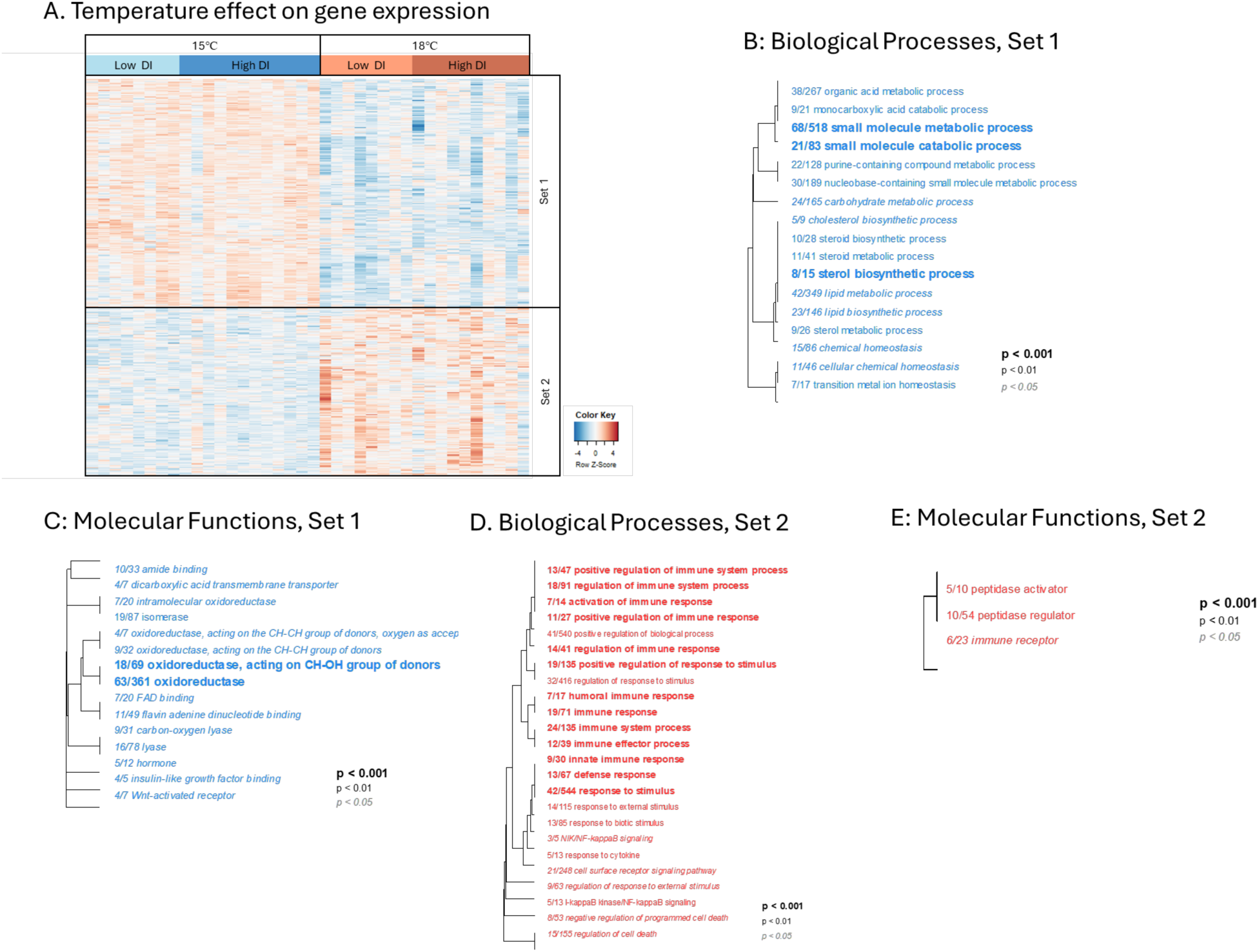
Rearing temperature effect on liver gene expression. A) Heatmap of expression patterns (log2 transformed) for genes that were significantly differentially expressed between 15°C or 18°C rearing environments (red and blue colors of cells represent higher and lower expression, respectively). Each column represents an individual, grouped by temperature then DI within temperature. Rows are genes with log2 transformed expression levels (color) and z-score normalized. In this analysis, Set-1 and Set-2 genes were more lowly or highly expressed, respectively, in 18°C compared to 15°C fish. GO terms under Biological Processes (B, D) and Molecular Functions (C, E) were functionally enriched in both Set1 (B, C) and Set2 genes (D, E). Dendrograms (B-E) represent hierarchical clustering of GO categories based on shared genes, which collapses redundant GO categories under the name of the lower-level (more specific) term. The fractions correspond to the number of genes that exceeded the unadjusted p < 0.05 over the total number of genes in that GO category.

We detected 357 DEGs that distinguished low from high DI fish, with 253 genes more highly expressed in high DI fish (Set-2, Figure 4A) and 104 genes more lowly expressed in high DI fish (Set-1). For Set-1 genes there was no functional enrichment for any GO categories. Set-2 genes were enriched for Biological Processes GO terms including, *lipid metabolic process*, *steroid and lipid biosynthetic process,* and *lipid transport* (Figure 4B). We observed few genes (26 genes) with expression showing a rearing temperature by DI interaction (Figure S3). These 26 genes were not enriched for any GO categories.

**Figure 4.**
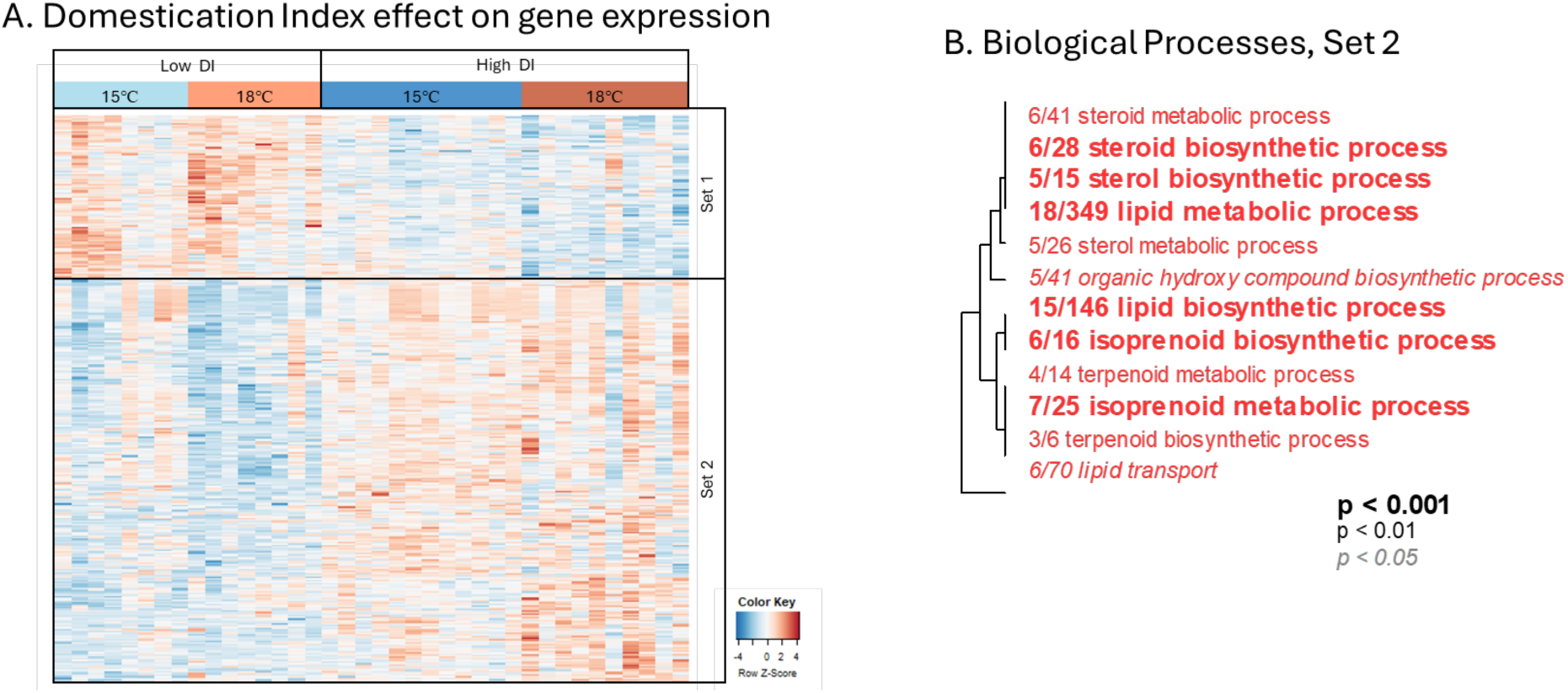
Domestication Index (DI) effect on liver gene expression. A) Heatmap of expression patterns for genes that were significantly differentially expressed between high and low DI fish (red and blue colors of cells represent higher and lower expression, respectively). Each column represents an individual, grouped by DI then temperature within DI. Rows are genes with log2 transformed expression levels (color) and z-score normalized. In this analysis, Set-1 and Set-2 genes were more lowly or more highly expressed, respectively, in high compared to low DI fish. GO terms under Biological Processes were functionally enriched for Set-2 genes only (B). The dendrogram represents hierarchical clustering of GO categories based on shared genes, which collapses redundant GO categories under the name of the lower-level (more specific) term. The fractions correspond to the number of genes that exceeded the unadjusted p < 0.05 over the total number of genes in that GO category.

### Differentially methylated regions

The filtered methylome dataset from liver tissue contained 7,259,297 CpG sites, which is 93.6% of all possible sites. Between rearing temperature treatments, we detected 1,385 DMRs, with 41 % of them hypermethylated (Set-2) and 59% hypomethylated (Set-1) in 18°C compared to 15°C fish (Figure 5). The mean size of DMRs was ∼620 bp. Fish from the 18°C treatment had on average 6% (range 4 to 15%) more CpG sites in the hypermethylated regions, and 10% (range 18 to 4%) fewer in the hypomethylated regions, compared to 15°C fish. Temperature DMRs were primarily enriched in intron regions (Table S1 and S2).

**Figure 5.**
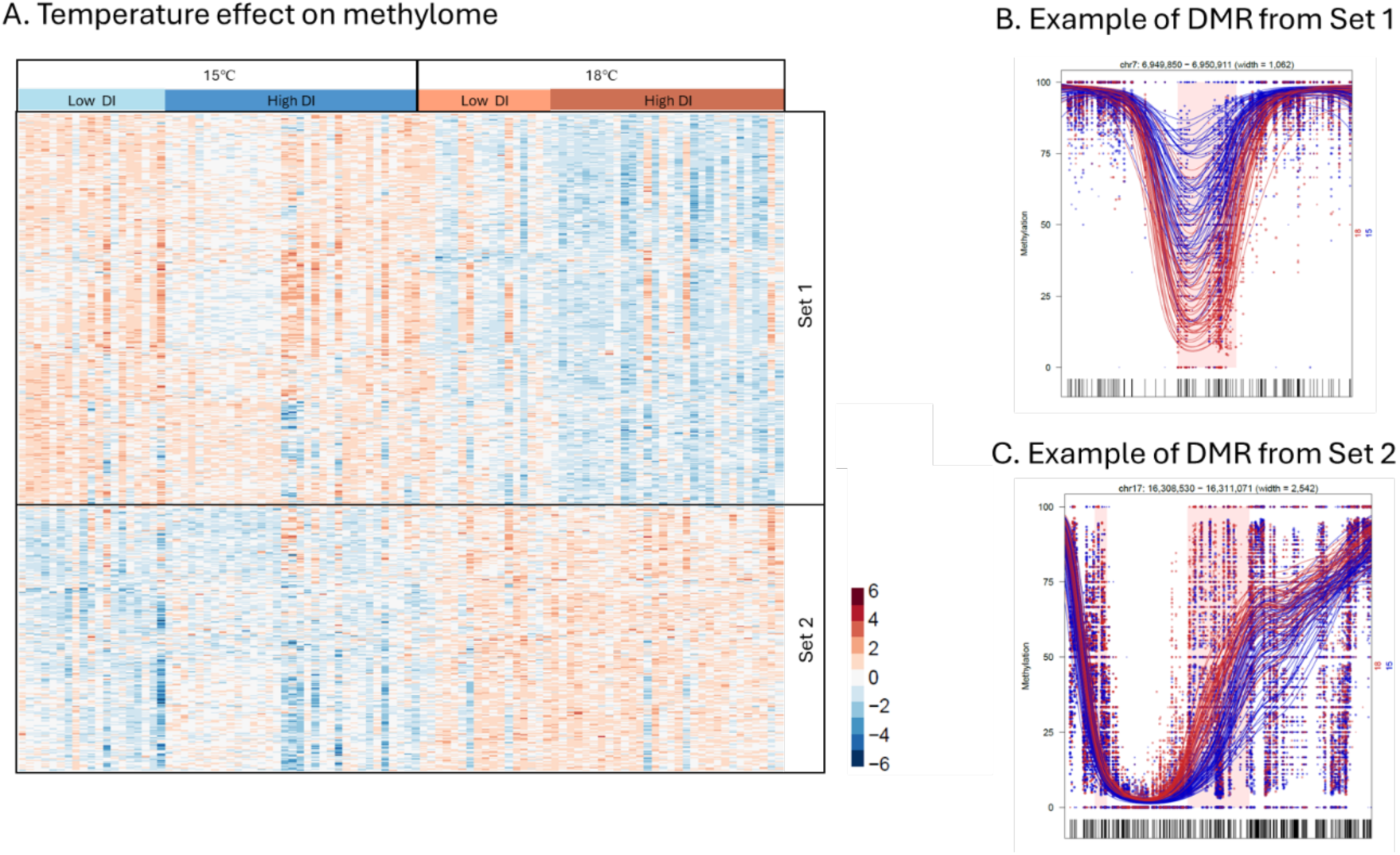
Rearing temperature effect on liver methylome. A) Heatmap of methylation patterns for differential methylation (DMRs) due to rearing temperature (red and blue colors of cells represent higher or lower percent methylation, respectively) (A). Each column represents an individual, grouped by temperature then by DI within temperature. Rows are genomic regions where percent methylation is z-score normalized. In this analysis, Set-1 and Set-2 are DMRs that were hypomethylated or hypermethylated, respectively, at 18°C compared to 15°C. Panels B and C show representative DMRs that were hypomethylated or hypermethylated, respectively, in 18°C compared to 15°C fish. The X-axis is the genomic position and Y-axis is the percent methylation within the DMR. Each dot represents the percent methylation value for a CpG site. Each line is a smoothed methylation value for each individual, color coded by treatment group: blue for 15°C and red for 18°C. The highlighted pink area is the region of differential methylation.

We detected 2,074 DMRs that distinguished high from low DI fish, with 63% of regions hypomethylated and 37% hypermethylated in high compared to low DI fish (Figure 6). The mean size of DMRs was ∼777 bp. Fish from the high DI group had on average 9% (range 4 to 32%) fewer CpG sites in the hypomethylated regions, and 10% (range 38 to 4%) more in the hypermethylated regions, compared to low DI fish. DI DMRs were primarily enriched in intergenic regions (Table S3 and S4). We did not detect any DMRs that had a temperature response that differed between low and high DI fish (e.g., no putative interactions between DI and temperature effects; Figure S4).

**Figure 6.**
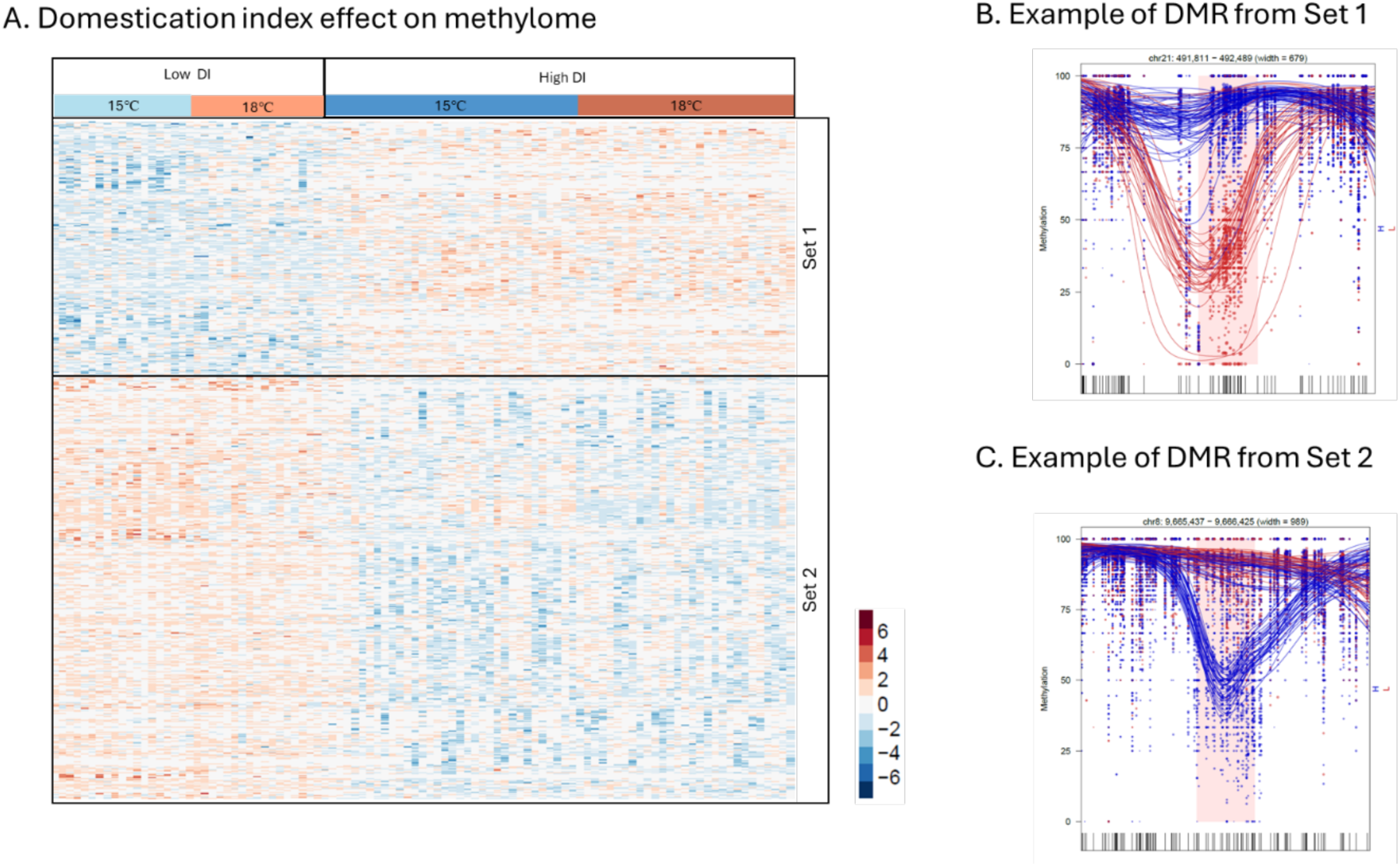
Domestication index effect on liver methylome. Heatmap of methylation patterns for differential methylation (DMRs) between low and high DI fish (red and blue colors of cells represent higher or lower percent methylation, respectively) (A). Each column represents an individual, grouped by DI then rearing temperature within DI. Rows are genomic regions where percent methylation is z-score normalized. In this analysis, Set-1 and Set-2 are DMRs that were hypermethylated or hypomethylated, respectively, in high compared to low DI fish. Panels B and C show representative DMRs that were hypermethylated or hypomethylated, respectively, in high compared to low DI fish. The X-axis is the genomic position and Y-axis is the percent methylation within the DMR. Each dot represents the percent methylation value for a CpG site. Each line is a smoothed methylation value for each individual, color coded by treatment group: blue for high DI fish and red for low DI fish. The highlighted pink area is the region of differential methylation.

We tested whether DMRs that varied between the livers of low and high DI fish identified in our study overlapped with those DMRs that varied across fin tissues in the first few generations of captivity identified by Habibi et al. (2024). Habibi et al. (2024) found 128, 122, and 180 DMRs between generation 0 and generation 1, 2, and 3 respectively. In contrast, we observed a much larger number of DMRs (2074) between low (∼5-6 generations) and high (∼ 9-10 generations) domestication categories. Of these, only 4, 10, and 21 DMRs overlapped with DMRs observed between generation 0 and generation 1, 2, and 3, respectively (Figure S5).

Finally, we investigated whether DMRs due to DI (low vs high DI from our study) and early captive rearing (generations 1 – 3 from Habibi et al. 2024) are regions of the genome that were also affected by rearing temperature. Using both DMR datasets from our study, we identified 130 common DMRs that were both affected by rearing temperature and variable between low and high DI fish, which was not more than expected by chance (p-value = 1). We found 8, 11, and 9 rearing-temperature induced DMRs overlapped with DMRs observed between generation 0 and generation 1, 2, and 3, respectively from Habibi et al. 2024 (Figure S6).

### Correlation of DMRs and DEGs

There was little overlap in the sets of genes that were both differentially expressed and differentially methylated, and even fewer showed significant correlation between differential expression and differential methylation. For the main effect of temperature, differential methylation and expression was detected for 89 genes, which was not more than expected by chance (p-value = 1). Percent methylation in a DMR was significantly correlated with gene expression for only 12 out of the 89 genes (Table S5), where correlation was positive and negative for 9 and 3 genes, respectively (Table S5). These genes were associated with promoters, introns, and exons in DMRs, but the functional region of the DMR (promotor, intron, exon) was not associated with whether the direction of differential methylation (hyper or hypo) had a positive or negative correlation with expression level (Table S5).

For the main effect of DI, there were 32 genes that were both differentially methylated and differentially expressed, which was not more than expected by chance (p-value = 0.99). Percent methylation in a DMR was significantly correlated with gene expression for 11 out of the 89 genes (Table S6), where all 11 had a negative correlation and were associated with DMRs found in promoters, introns, and exons (Table S6).

### Genetic differentiation between low and high DI fish

We estimated genome-wide genetic differentiation (F_ST_) between low and high DI adult fish. Genome-wide average F_ST_ was 0.008, which was higher than expected by chance (p-value =0; all permutated F_ST_ values were below 0.008). In addition, some regions of the genome displayed elevated F _ST_ around 0.2 and there were 0.3% of SNPs that had an F_ST_ greater than 0.1 (Figure S7).

## Discussion

Hatchery environments can quickly induce shifts in physiology within and across generations that may be of consequence for fish performance. We tested whether rearing temperatures affected upper thermal tolerance (acclimation) and whether thermal physiology has been changing in hatchery fish (evolution or imprinting), and examined the underlying molecular mechanisms. Our key findings were: 1) Rearing at elevated temperature increased thermal tolerance limits of Delta smelt, consistent with adaptive acclimation. This exposure to elevated temperature altered genome-wide gene expression and DNA methylation in the liver; 2) Delta smelt with a longer history of hatchery domestication had higher upper thermal tolerances, consistent with heritable modification of thermal physiology, accompanied by extensive differentiation in liver transcriptomes and methylomes; 3) Delta smelt with high and low domestication levels in this study did not differ in their molecular plastic response to temperature. That is, there were no DI-by-temperature interactions for transcriptomic nor methylomic data. While we also did not observe a higher level interaction between DI and temperature in our CTMax data, more nuanced family-level analysis indicated that families with stronger temperature acclimation abilities (steeper CTMax reaction norm between 15°C and 18°C rearing treatments) were more common within low DI fish, whereas families with more shallow thermal reaction norms were more common within high DI fish. This is consistent with a subtle reduction in temperature acclimation plasticity that accompanied domestication. We conclude that domestication has resulted in higher thermal tolerance, but perhaps at the cost of reduced thermal plasticity.

### Rearing temperature effects on physiology, transcriptome, and methylome

Delta smelt displayed a modest but statistically significant ability to acclimate to different thermal environments (thermal plasticity). We observed a small increase in CTMax (0.63°C) for fish reared and acclimated at 18°C compared to 15°C (Figure 1). This contrasts with other findings where juvenile and adult Delta smelt showed no thermal acclimation across a similar temperature range (Komoroske et al. 2014). Only when adult Delta smelt have been acclimated to potentially stressful temperatures, such as 20°C, are increases in CTMax observed (Davis et al. 2019a). This suggests that thermal acclimation abilities vary between life stages in Delta smelt, where earlier life stages are more plastic and sensitive to smaller increases in temperature. However, since a three-degree difference in rearing environment shifted CTMax by only 0.63 degrees (Figure 1A), persistence of Delta smelt in warming environments will likely depend on acclimation plus other phenomena such as evolutionary change.

Despite modest gains in thermal tolerance, rearing temperature had substantial genome-wide effects on methylation and gene expression. Rearing temperature caused differential expression of 1,609 genes (Figure 3). Genes that were upregulated at higher rearing temperature were primarily involved in immune functions, a common response to prolonged exposure to elevated temperatures in fish (Huang et al. 2018; Zhang et al. 2021). Upregulation of immune genes is a core component of the generalized stress response and is utilized in the heat stress response as well (Li et al. 2019). Rearing temperature also affected transcription of genes related to metabolic processes and oxidative stress response (i.e., oxidoreductase activity). This response may reflect physiological changes required to compensate for elevated metabolic demands, and associated production of reactive oxygen species (ROS) at elevated temperature (Person-Le Ruyet et al. 2004; Vergauwen et al. 2010; Madeira et al. 2013).

Different rearing temperatures also resulted in genome wide change in the methylome (1,385 DMRs; Figure 7) as is observed in other fish species (Skjærven et al. 2014; Anastasiadi et al. 2017). Environmentally induced epigenetic changes can modulate gene expression and thereby contribute to environmentally induced phenotypic changes (Vineis et al. 2017). For example, promoter methylation may repress transcription (Boyes and Bird 1992; Jones and Takai 2001), whereas gene-body methylation may promote transcriptional stability and alternative splicing (Maunakea et al. 2010). However, we found little overlap of DMRs and DEGs in response to temperature, and little overlap in the associated molecular pathways and functions implicated by DMRs and DEGs (GO enrichment). Therefore, the mechanistic connection between differential methylation and differential gene expression is not obvious in our study. A similar lack of overlap is commonly observed, including in other aquatic species (Skjærven et al. 2018; Anastasiadi et al. 2021; Jones and Griffitt 2022; Bogan 2024). Of course, methylation is not the only molecular mechanism through which environmental change is transduced into transcriptional change, and those mechanisms (e.g., regulation of transcription factors, histone modifications, etc.) may be more functionally related to the transcriptional changes that we observe. Furthermore, it is plausible that this lack of overlap reflects the different temporal dynamics of environmentally-induced methylation and transcription. Perhaps temperature-induced methylation induces differential transcription, where the methylation change is stable but the transcriptional change is transient. Indeed, methylome changes induced by temperature during early life can persist for multiple years (Anastasiadi et al. 2021). It is also plausible that temperature-induced methylation serves to stabilize and constrain temperature induced changes in transcription. The functional connection between temperature induced methylation and transcription underlying thermal acclimation clearly merits further study, where experiments that illuminate the temporal dynamics that connect temperature change with methylation and transcription, and ultimately physiology, would be particularly valuable.

### Domestication effects on physiology, transcriptome, and methylome

Highly domesticated (high DI) fish displayed higher CTMax compared to low DI fish at both rearing temperatures, suggesting that thermal physiology has changed as a consequence of domestication. The CTMax for high DI fish was 1°C higher than for low DI fish, which represents a larger increase in CTMax than that caused by thermal acclimation. Other studies have documented either no differences in CTMax or lower CTMax in domesticated fish compared to their wild counterparts in trout, salmon, and minnow (Carline and Machung 2001; Chen et al. 2015; Hirakawa and Salinas 2020).

There was also a domestication effect on the transcriptome and methylome. We observed 356 DEGs that distinguished low from high DI fish (Figure 4). These genes were functionally enriched in metabolic and biosynthetic processes that were more highly expressed in higher DI fish. Similar functional enrichment was observed among genes that distinguished hatchery fish from their wild counterparts in Steelhead trout and Nile tilapia (Christie et al. 2016; Konstantinidis et al. 2020), suggesting that divergence in metabolic and biosynthetic traits may be generally associated with hatchery environments for multiple species. It is plausible that differential expression of genes involved in metabolic and biosynthetic processes underlie the differences in thermal physiology that we observed between low and high DI fish. However, CTMax is the only physiological trait that we measured; it is likely that some of the transcriptome differences that we detected may underlie physiological traits that we did not measure but that also differ between low and high DI fish. We conclude that the hatchery has imposed an environmental change that has manifested as physiological and molecular changes that accumulate and drive divergence across generations.

Domestication appears to have caused a change in thermal physiology limits, as well as divergence in the plasticity of thermal tolerance (acclimation ability) in this study. At a broad genotype level (i.e., DI), there was no statistically significant genotype-by-environment interaction between DI and rearing temperature for CTMax. However, grouping individuals into families and comparison across families allows for greater nuance in examining heritable genetic variation in thermal physiology traits. Since each family was split between the two rearing environments, we could measure average CTMax for related individuals reared in the two environmental conditions and draw a reaction norm for each family representing the rearing temperature effect on CTMax (thermal plasticity; Figure 1b). After comparing the variation in slopes of reaction norms among families within and between DI groups we detected a subtle but statistically significant relationship between domestication and thermal plasticity. Families with high plasticity were more common among the low DI fish, whereas families with lower plasticity were enriched among the high DI fish (Figure 1c). This is consistent with domestication favoring higher CTMax, but perhaps at the cost of reduced thermal plasticity. This could be a consequence of hatchery environments being more thermally stable than natural environments in which high plasticity could be favored (Waddington 1959; Crispo 2007). At the transcriptomic level, we found minimal evidence for DI-by-temperature interactions (only 25 genes). Future transcriptomics studies designed with sampling to capture the effect of family could offer insight into molecular mechanisms underlying domestication impacts on thermal plasticity.

Traits that have diverged with hatchery domestication could be a consequence of evolutionary (genetic) change or inherited but non-genetic (e.g., epigenetic) change. Multiple traits in addition to thermal physiology and transcription appear to have been diverging in the Delta smelt refuge hatchery, including reproductive success and maturating timing (Finger et al. 2018; LaCava et al. 2023; Tsai et al. 2023). What are the mechanisms that explain this multi-generational divergence? It is clear that there is much epigenetic variation that distinguishes high from low DI fish (over 2,000 DMRs). It is plausible that the novel hatchery environment induced epigenetic changes, and that these were stably inherited within hatchery lineages across generations (epigenetic imprinting). Alternatively, genetic change (caused by random drift or selection) could have evolved to distinguish low from high DI fish, with epigenetic change as a consequence (e.g., (Silliman et al. 2023). If the former is true, one may predict that 1) epigenetic changes induced in wild fish introduced into the new hatchery environment would be stably inherited in subsequent generations, and 2) that features of hatchery environments (like temperature) would induce epigenetic change that is stably inherited. However, the evidence so far does not support these predictions. There is no overlap between temperature-induced methylome change and methylome changes that distinguish high from low DI fish (for DMRs, no interaction between temperature and DI). Furthermore, epigenetic changes were not stably inherited across the first few generations of hatchery fish (Habibi et al. 2024), nor were DMRs consistent between early (Habibi et al. 2024) and later-generation (this study) domesticated lineages (Figure S5). However, we note that differences in experimental design and tissue types may limit inferences and conclusions about the similarities and differences in the outcomes of the two experiments. Finally, if DMRs were responsible for diverged thermal physiology and other traits we might expect functional enrichment of related genetic pathways. We found no functional GO enrichment among DMRs that distinguish high from low DI fish. We conclude that there is limited evidence that inherited epigenetic imprinting contributes to the divergence of traits observed in domesticated lineages of hatchery-reared Delta smelt.

To account for domestication divergence, an alternative explanation to epigenetic imprinting is evolutionary change. If evolutionary change is responsible, then one would predict genetic differentiation between high and low DI fish. Indeed, we find that genetic changes have accumulated that differentiate high from low DI fish: genome-wide average F_ST_ between these two groups was 0.008 indicating a subtle influence of random-neutral genetic drift. Furthermore, few regions of the genome have accumulated relatively high F_ST_ (e.g., >0.1) between low and high DI fish suggesting the influence of natural selection (Figure S7). Even though the refuge hatchery has been carefully genetically managed, genetic divergence seems to still have accumulated. But it is not clear whether drift or selection has driven the divergence in traits that we and others have observed in highly domesticated fish. If natural selection has driven elevated CTMax in high DI fish, one might predict reduced inter-individual variation in CTMax and/or gene expression among high DI fish compared to low DI fish. We do not observe this (Figure 1A, 2). Furthermore, it is not clear that hatchery temperatures are sufficiently different from the wild that they would affect fitness. Instead, CTMax variation may be evolving by neutral drift. Or CTMax variation may be linked to other traits, such as those that are evolving by natural selection to support the higher average stress tolerance and fitness of high compared to low DI fish in the hatchery environment (Finger 2018, LaCava 2023). Ongoing studies are using quantitative genetic approaches to identify genetic variants that are associated with inter-individual variation in CTMax. One could then test whether those loci show patterns of variation over time that are consistent with selection or drift.

### Conclusion

Given the rising temperatures in the San Francisco estuary, elevated thermal tolerance that is resilient and sustainable may be necessary for Delta smelt persistence. Experimental releases of Delta smelt into the wild from the hatchery are already underway (U.S. Fish and Wildlife Service 2020) and hatchery practices that can increase the thermal tolerance, either through warm acclimation or selection of fish from families with superior high-temperature performance, may be crucial for ensuring long-term survival of the species. Our results suggest that rearing at elevated temperatures can achieve modest increases in thermal tolerance, and that more highly domesticated fish have evolved additional gains in high temperature tolerance. This may be a rare case of unintentional domestication selection that increases performance (high temperature tolerance) in the same direction as that needed to persist in changing natural environments. But this conclusion should be treated with caution since domestication selection may have altered thermal plasticity and other traits in directions that may be less favorable upon reintroduction to the wild.

## Acknowledgements

This research was supported by the California Department of Fish and Wildlife (Prop 1: Watershed Restoration Grant Program / Delta Water Quality and Ecosystem Restoration Program. Grant Q1996041, PIs Dr. Andrew Whitehead, Dr. Nann Fangue, Dr. Tien-Chieh Hung). We thank current and former FCCL staff for fish maintenance (especially Luke Ellison and William Mulvaney). We thank UC Davis staff that helped with physiological measurements (Jenn Roach, Dr. Nicole McNabb, Ashley De La Torre, and Anthony Tercero), and Dr. Brittany Davis for help with CTMax set up and analysis. The sequencing was carried out at the UC Davis Genome Center DNA Technologies and Expression Analysis Core, supported by NIH Shared Instrumentation Grant 1S10OD010786-01. We acknowledge the High Performance Computing Core Facility at the University of California, Davis, for providing computational resources that have contributed to the research results reported in this paper.

## Data Availability

Sequencing data are deposited in the National Center for Biotechnology Information’s Short Reads Archive (RNASeq: BioProject PRJNA1112349; WGBS: BioProject PRJNA1117032). Scripts and code for RNASeq and WGBS analyses are available on GitHub https://github.com/WhiteheadLab/Delta_Smelt_Epigenetics_Transciptomics_copy.

## Notes

### Competing Interest Statement

The authors have declared no competing interest.

### Summary of Updates

The author name Md Moshiur Rahman was originally spelled incorrectly. This mistake has been corrected in the revised version of the manuscript.

