## Supplemental Materials Figures S1-S7 Tables S1-S6 for "Highly and lowly domesticated endangered fish from a conservation hatchery diverge in their thermal physiology, transcriptome, and methylome"

**SUPPLEMENTAL FIGURES and TABLES**

### Epigenetic and transcriptomic responses to warming are similar between highly and lowly domesticated Delta smelt from a conservation hatchery

Joanna S. Griffiths[1,2], Amanda J. Finger[3], Melinda R. Baerwald[4], Md Moshiur Rahman[5], Tien-Chieh Hung[5], Nann A. Fangue[6*], and Andrew Whitehead[1*]

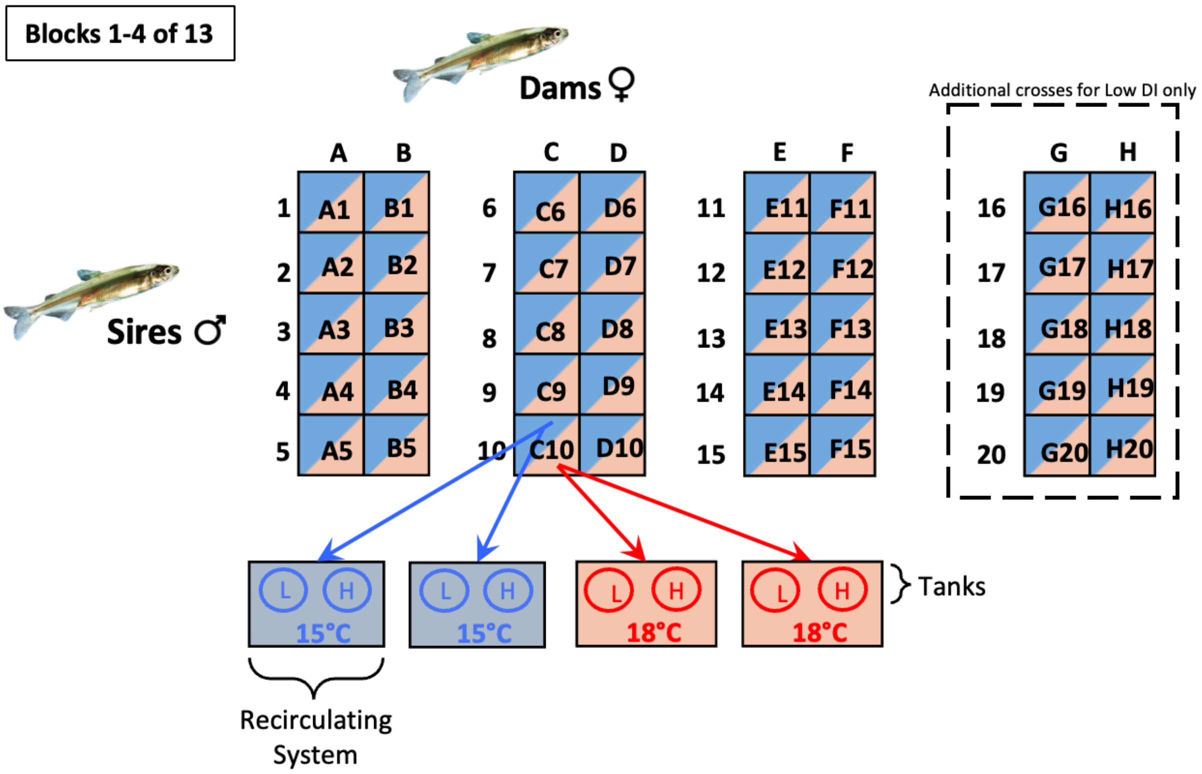


Figure S1: Factorial breeding block design and offspring rearing temperature experimental design. Families were generated from adult smelt of low or high DI category. Each block consisted of 5 males (numbers) crossed with 2 females (letters) from the same DI category. Each family was designated by the number and letter combo of its parents (i.e., C10). There were 4 blocks for the low DI group and 3 blocks for the high DI group. There were a total of 49 parents crossed and 70 families generated. Each family was evenly split into four groups and reared at either 15C or 18C (with replicates for each temperature). Fish from the same DI category (low or high) were kept together in a single tank, fed by a single header tank for each temperature and replicate (i.e., Closed recirculating "system").

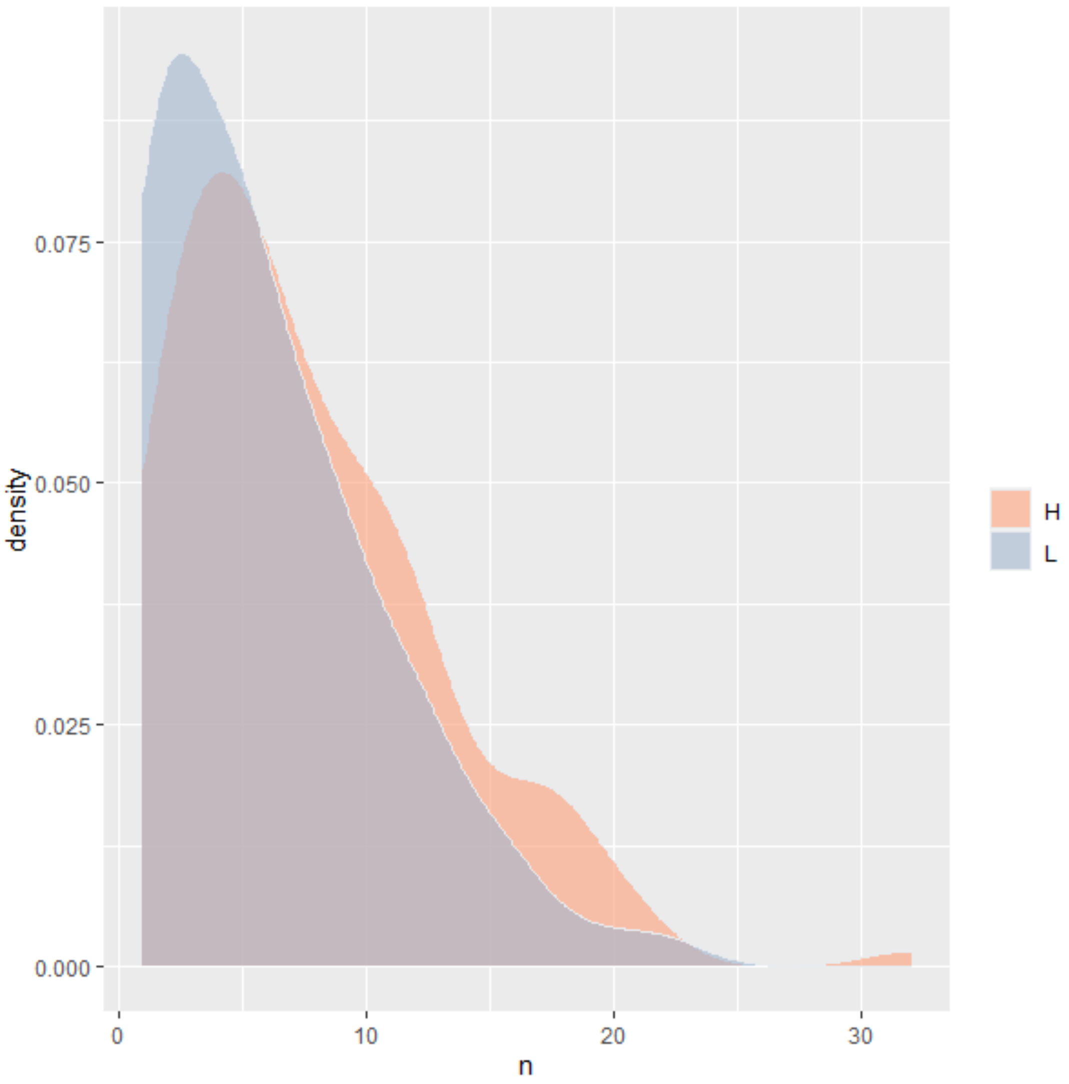


Figure S2: Density plot of the number of surviving offspring from families of low (blue) or high (orange) DI fish. Density plots are similar between the two groups of fish, confirming that survival between low and high DI families is similar.

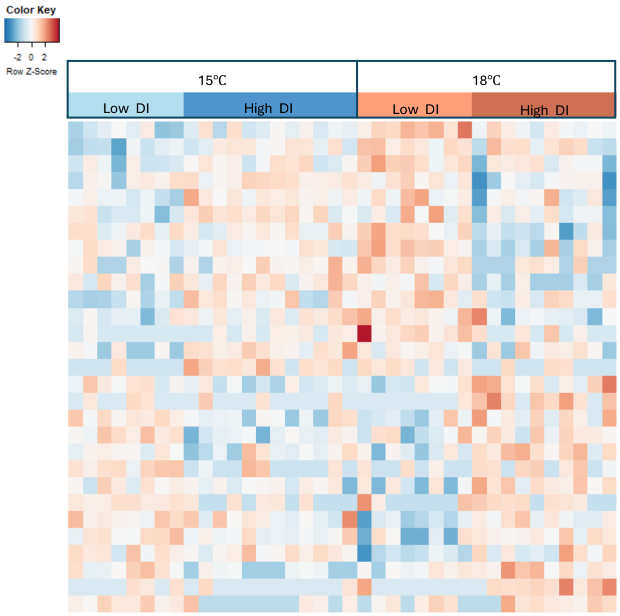


Figure S3: Temperature by Domestication Index (DI) interaction effects on gene expression (red colors represents higher expression and blue colors represents lower expression). Heatmap of expression patterns for genes that were significantly differentially expressed. Each column represents an individual from either a high or low DI background, reared at either 15°C or 18°C . Rows are sorted by gene expression level and each row is z-score normalized.

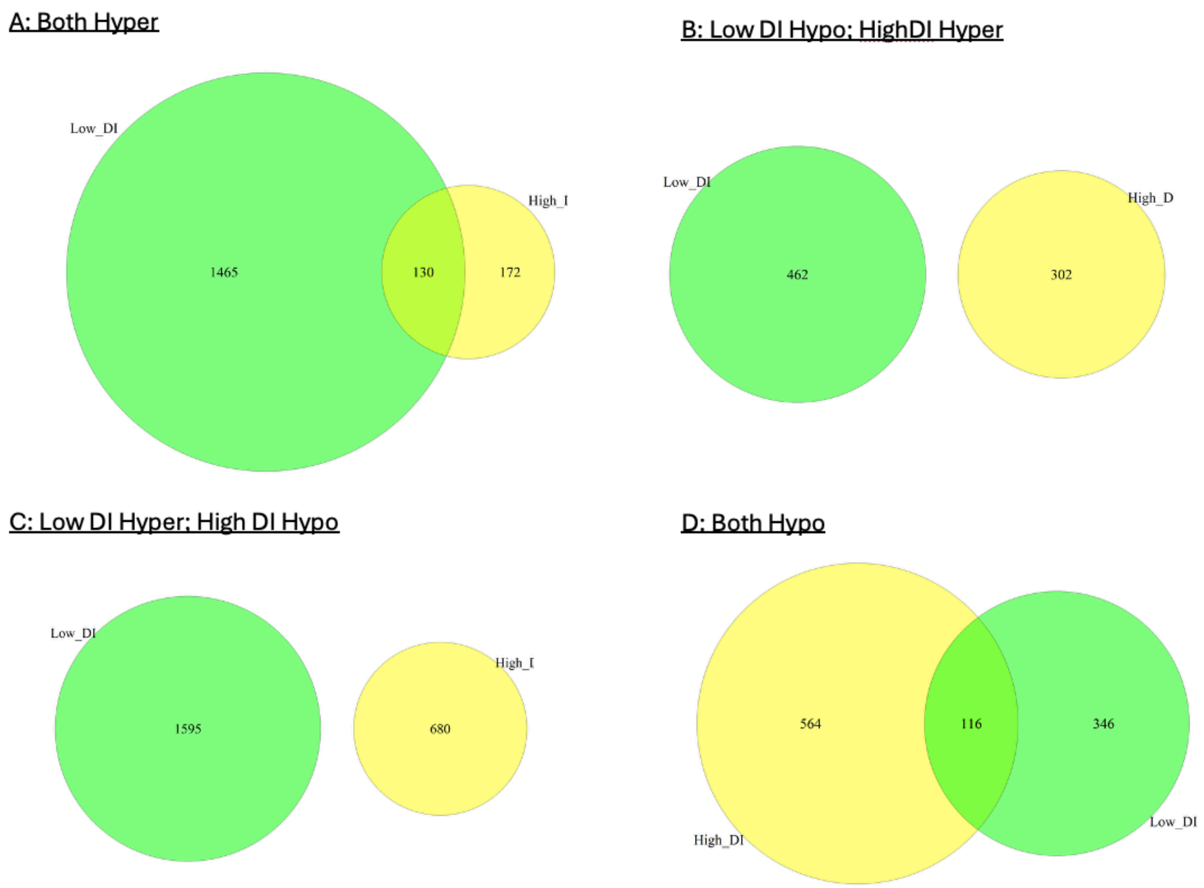


Figure S4: Venn diagram of DMRs due to acclimation temperature subsetted for low DI fish only and high DI fish only. We then identified DMRs that were hypermethylated in both datasets (A), hypomethylated in low DI fish and hypermethylated in high DI fish (B), hypermethylated in low DI fish, hypomethylated in high DI fish (C), and hypomethylated in both datasets. An interaction effect of acclimation temperature and DI was defined as any DMRs that were hypermethylated in one group and hypomethylated in the other group. Using this approach, we did not find any overlap of DMRs that were hypermethylated in one group and hypomethylated in the other group (B and C).

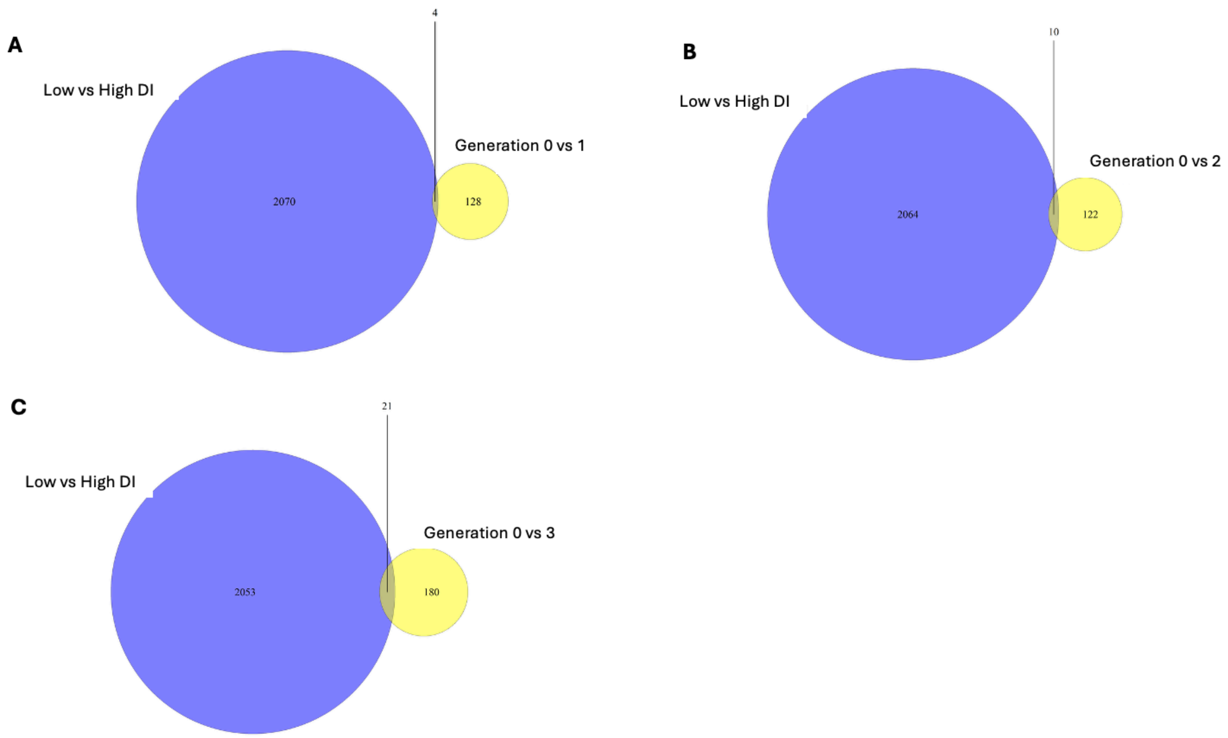


Figure S5: Venn diagram of DMRs due to domestication index found in our study (blue) compared to DMRs found in Habibi et al. 2024 (yellow). Overlap of DMRs between low and high DI fish and those between generation 0 to generation 1 from Habibi et al. 2024 (A). Overlap of DMRs between low and high DI fish and those between generation 0 to generation 2 from Habibi et al. 2024 (B). Overlap of DMRs between low and high DI fish and those between generation 0 to generation 3 from Habibi et al. 2024 (C).

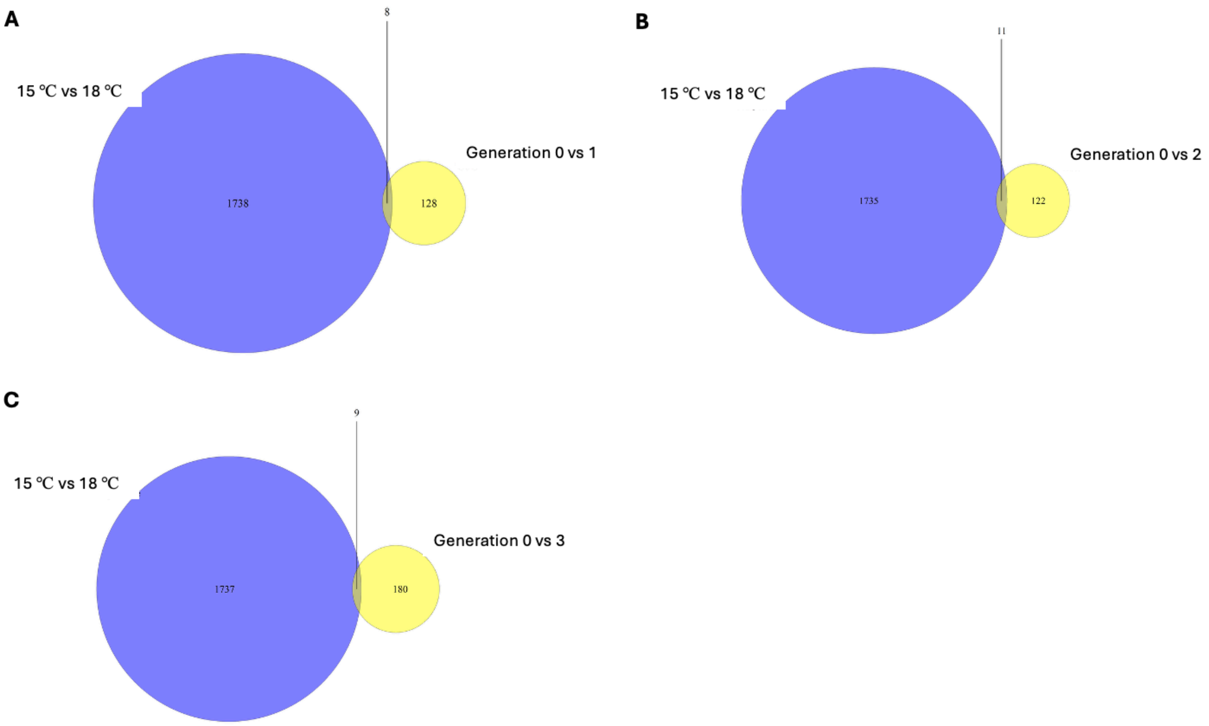


Figure S6: Venn diagram of DMRs due to rearing temperature (blue) found in our study compared to DMRs found in Habibi et al. 2024 (yellow). Overlap of DMRs between fish reared at 15 ℃ vs 18 ℃ and those between generation 0 to generation 1 from Habibi et al. 2024 (A). Overlap of DMRs between fish reared at 15 ℃ vs 18 ℃ and those between generation 0 to generation 2 from Habibi et al. 2024 (B). Overlap of DMRs between fish reared at 15 ℃ vs 18 ℃ and those between generation 0 to generation 3 from Habibi et al. 2024 (C).

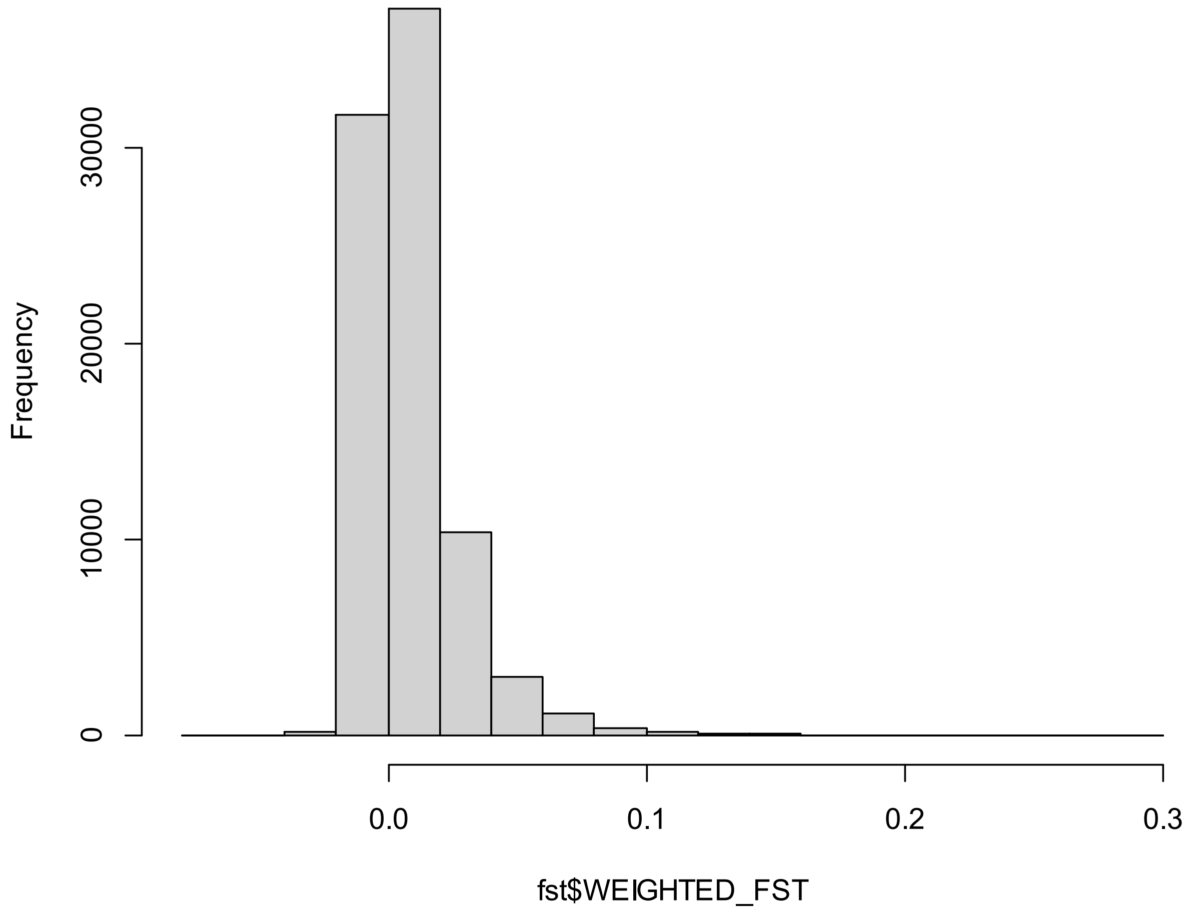


Figure S7: Histogram of Weir pairwise $F_{ST}$ (window size of 10,000 bp and a sliding window of 5000 bp) between low and high DI progenitors.

Table S1: HOMER annotation of hypermethylated DMRs due to acclimation temperature. DMRs are annotated based on overlap to TTS (transcription termination site), exons, introns, intergenic, or promoters. The relative enrichment of DMRs in each set of genomic annotations is displayed along with log pvalue (i.e. enrichment of the peaks in promoter regions, exons, etc.)

| **Annotation** | **Number of DMRs** | **Log2 Ratio (obs/exp)** | **LogP enrichment (+values depleted)** |
|---|---|---|---|
| TTS | 38 | 0.015 | -0.7 |
| Exon | 64 | -0.544 | 7.778 |
| Intron | 399 | 0.279 | -15.982 |
| Intergenic | 181 | -0.281 | 7.11 |
| Promoter | 47 | -0.073 | 0.93 |

Table S2: HOMER annotation of hypomethylated DMRs due to acclimation temperature. DMRs are annotated based on overlap to TTS (transcription termination site), exons, introns, intergenic, or promoters. The relative enrichment of DMRs in each set of genomic annotations is displayed along with log pvalue (i.e. enrichment of the peaks in promoter regions, exons, etc.)

| **Annotation** | **Number of DMRs** | **Log2 Ratio (obs/exp)** | **LogP enrichment (+values depleted)** |
|---|---|---|---|
| TTS | 61 | 0.218 | -2.056 |
| Exon | 91 | -0.516 | 9.503 |
| Intron | 575 | 0.326 | -29.405 |
| Intergenic | 222 | -0.467 | 20.317 |
| Promoter | 68 | -0.021 | 0.727 |

Table S3: HOMER annotation of hypermethylated DMRs due to domestication index (DI). DMRs are annotated based on overlap to TTS (transcription termination site), exons, introns, intergenic, or promoters. The relative enrichment of DMRs in each set of genomic annotations is displayed along with log pvalue (i.e. enrichment of the peaks in promoter regions, exons, etc.)

| Annotation | Number of DMRs | Log2 Ratio (obs/exp) | LogP enrichment (+values depleted) |
|---|---|---|---|
| TTS | 58 | 0.548 | -5.837 |
| Exon | 61 | -0.69 | 11.342 |
| Intron | 218 | -0.67 | 48.302 |
| Intergenic | 347 | 0.581 | -40.834 |
| Promoter | 85 | 0.705 | -11.673 |

Table S4: HOMER annotation of hypomethylated DMRs due to domestication index (DI). DMRs are annotated based on overlap to TTS (transcription termination site), exons, introns, intergenic, or promoters. The relative enrichment of DMRs in each set of genomic annotations is displayed along with log pvalue (i.e. enrichment of the peaks in promoter regions, exons, etc.)

| Annotation | Number of DMRs | Log2 Ratio (obs/exp) | LogP enrichment (+values depleted) |
|---|---|---|---|
| TTS | 98 | 0.542 | -8.634 |
| Exon | 110 | -0.602 | 14.743 |
| Intron | 415 | -0.504 | 50.965 |
| Intergenic | 606 | 0.622 | -79.143 |
| Promoter | 76 | -0.22 | 2.399 |

Table S5: Genes that showed a significant correlation (with Bonferroni correction applied) between expression levels and percent methylation for DEG and DMRs due to acclimation temperature. Transcript ID, Gene ID, Gene product, and GO ID are described. We also note where the DMR region was located (intron, exon, or promoter region for each gene), whether the correlation was negative or positive. Finally we note the pvalue and adjusted r-2 results of the linear model test for each gene.

| Transcript ID | Gene ID | Gene product | GO ID/function | DMR location | Correlation | P-value | Adjusted r-2 |
|---|---|---|---|---|---|---|---|
| XM_047020039.1 | apln | apelin | No GO ID | intron | neg | 2.823004e-05 | 0.381 |
| XM_047020579.1 | mogat2 | monoacylglycerol O-acyltransferase 2 | No GO ID | exon | pos | 3.036420e-06 | 0.4528 |
| XM_047020975.1 | hpda | 4-hydroxyphenylpyruvate dioxygenase a | No GO ID | intron | neg | 2.833100e-06 | 0.4549 |
| XM_047022316.1 | LOC124469183 | LRP2 binding protein | No GO ID | promoter | pos | 3.880637e-09 | 0.6231 |
| XM_047023175.1 | lrrc17 | leucine rich repeat containing 17 | No GO ID | intron | neg | 3.634301e-07 | 0.5138 |
| XM_047022638.1 | cog5 | component of oligomeric golgi complex 5 | No GO ID | intron | neg | 3.549403e-04 | 0.2892 |
| XM_047027751.1 | LOC124472742 | uncharacterized | No GO ID | exon | neg | 4.305805e-06 | 0.4421 |
| XM_047039291.1 | LOC124480190 | WAP four-disulfide core domain protein 2-like | No GO ID | promoter | neg | 2.384565e-04 | 0.3044 |
| XM_047040555.1 | LOC124480897 | ATPase family AAA domain-containing protein 3-like | No GO ID | exon | neg | 6.063330e-05 | 0.3545 |
| XM_047044466.1 | serping1 | serpin peptidase inhibitor, clade G (C1 inhibitor), member 1 | No GO ID | exon | neg | 1.347628e-04 | 0.3256 |
| XM_047044619.1 | rhof | ras homolog family member F | No GO ID | intron | neg | 1.353223e-04 | 0.3255 |
| XR_006957121.1 | LOC124477337 | uncharacterized | No GO ID | exon | pos | 6.602525e-05 | 0.3515 |

Table S6: Genes that showed a significant correlation (with Bonferroni correction applied) between expression levels and percent methylation for DEG and DMRs due to domestication index (DI). Transcript ID, Gene ID, Gene product, and GO ID are described. We also note where the DMR region was located (intron, exon, or promoter region for each gene), whether the correlation was negative or positive. Finally we note the  pvalue and adjusted r-2 results of the linear model test for each gene.

| Transcript ID | Gene ID | Gene product | GO ID/function | DMR location | Correlation | P-value | Adjusted r-2 |
|---|---|---|---|---|---|---|---|
| XM_047020528.1 | arr3a | arrestin 3a, retinal (X-arrestin) | No GO ID | intron | neg | 1.312e-06 | 0.4778 |
| XM_047025846.1 | LOC124471360 | zinc finger and SCAN domain-containing protein 2-like | No GO ID | exon | neg | 3.854e-05 | 0.3703 |
| XM_047028340.1 | LOC124473082 | beta-1,3-galactosyltransferase 2-like | No GO ID | intron | neg | 0.0002076 | 0.3096 |
| XM_047032478.1 | LOC124475690 | type-2 ice-structuring protein-like | No GO ID | promoter | neg | 0.0001354 | 0.3255 |
| XM_047037577.1 | LOC124479076 | galectin-3-binding protein A-like | No GO ID | intron | neg | 7.447e-08 | 0.5551 |
| XM_047038435.1 | igf2bp1 | insulin-like growth factor 2 mRNA binding protein 1 | No GO ID | intron | neg | 6.419e-05 | 0.3525 |
| XM_047038932.1 | si:dkey-283b1.6 | uncharacterized | No GO ID | intron | neg | 4.635e-10 | 0.6657 |
| XM_047046573.1 | abca12 | ATP-binding cassette, sub-family A (ABC1), member 12 | No GO ID | exon | neg | 0.000298 | 0.2959 |
| XM_047046575.1 | abca12 | ATP-binding cassette, sub-family A (ABC1), member 12 | No GO ID | intron | neg | 1.679e-05 | 0.3985 |
| XM_047049358.1 | gba3 | glucosidase, beta, acid 3, transcript variant X1 | No GO ID | exon | neg | 5.467e-07 | 0.5026 |
| XR_006956573.1 | LOC124471944 | uncharacterized | No GO ID | promoter | neg | 2.825e-07 | 0.5206 |
